# Do Topographic Deep ANN Models of the Primate Ventral Stream Predict the Perceptual Effects of Direct IT Cortical Interventions?

**DOI:** 10.1101/2024.01.09.572970

**Authors:** Martin Schrimpf, Paul McGrath, Eshed Margalit, James J. DiCarlo

## Abstract

Ever-advancing artificial neural network (ANN) models of the ventral visual stream capture core object recognition behavior and the neural mechanisms underlying it with increasing precision. These models take images as input, propagate through simulated neural representations that resemble biological neural representations at all stages of the primate ventral stream, and produce simulated behavioral choices that resemble primate behavioral choices. We here extend this modeling approach to make and test predictions of neural intervention experiments. Specifically, we enable a new prediction regime for topographic deep ANN (TDANN) models of primate visual processing through the development of *perturbation modules* that translate micro-stimulation, optogenetic suppression, and muscimol suppression into changes in model *neural activity*. This unlocks the ability to predict the *behavioral* effects from particular neural perturbations. We compare these predictions with the key results from the primate IT perturbation experimental literature via a suite of nine corresponding benchmarks. Without any fitting to the benchmarks, we find that TDANN models generated via co-training with both a spatial correlation loss and a standard categorization task qualitatively predict all nine behavioral results. In contrast, TDANN models generated via random topography or via topographic unit arrangement after classification training predict less than half of those results. However, the models’ quantitative predictions are consistently misaligned with experimental data, over-predicting the magnitude of some behavioral effects and under-predicting others. None of the TDANN models were built with separate model hemispheres and thus, unsurprisingly, all fail to predict hemispheric-dependent effects. Taken together, these findings indicate that current topographic deep ANN models paired with perturbation modules are reasonable guides to predict the qualitative results of direct causal experiments in IT, but that improved TDANN models will be needed for precise quantitative predictions.

## Introduction

Some Artificial Neural Networks (ANNs) have recently been shown to be surprisingly good models of the neural mechanisms underlying a range of primate behaviors. This is especially true for models in the sensory domains vision (Yamins et al., 2014; Cadieu et al., 2014; Khaligh-Razavi and Kriegeskorte, 2014; Yamins and DiCarlo, 2016; Cadena et al., 2017; Schrimpf et al., 2018; Kubilius et al., 2019; Dapello et al., 2020; Schrimpf et al., 2020; Zhuang et al., 2021; Geiger et al., 2022; Dapello et al., 2023; Doerig et al., 2023), audition (Kell et al., 2018; Saddler et al., 2021), and somatosensation (Zhuang et al., 2017), but also for models of language processing (Schrimpf et al., 2021; Caucheteux and King, 2022; Toneva and Wehbe, 2019) and navigation (Nayebi et al., 2021). For a given domain, the general approach is to first show the exact same stimuli (e.g. images for the domain of vision) to both model “subjects” and to human or animal subjects. The candidate ANN model is then evaluated both on how closely its internal “neural” activity matches the neural activity measured experimentally in the subjects, and on how well its behavioral choices match the same types of behavioral choices measured experimentally from the subjects. For instance, the current leading ANN models of the ventral stream quite accurately predict image-by-image patterns of neural activity recorded from areas V1, V2, V4, and inferotemporal cortex (IT), as well as the core visual object recognition behaviors that the ventral stream supports within the larger domain of visual capabilities (Schrimpf et al., 2020).

However, a crucial area of neuroscience research has so far been neglected by these models: direct neural perturbations that causally link neural activity to behavioral outcomes. Such experimental studies perturb a piece of neural tissue and measure the effects on perception as measured via behavioral reports. Accurate predictions of the behavioral effects of neural interventions are a crucial stepping stone to the development of brain-machine-interfaces. In the primate visual ventral stream, the effect of neural perturbations in high-level visual area IT has been linked to changes in a number of perceptual reporting behaviors including face-related judgements (Afraz et al., 2006, 2015; Moeller et al., 2017; Sadagopan et al., 2017), orientation estimates (Adab and Vogels, 2016), glossiness estimates (Baba et al., 2021), and object category reports (Kawasaki and Sheinberg, 2008; Rajalingham and DiCarlo, 2019). Each of these studies uses one or more direct neural perturbation methods, including electrical micro-stimulation, optogenetic suppression, and muscimol suppression.

Efforts to develop machine executable models that attempt to explain and predict the behavioral effects of such perturbation experiments promise two benefits. First, such efforts will guide the development of new, more accurate scientific models. Specifically, these tests force causal links from e.g. model IT to behavioral report – current empirical benchmarking tests on the other hand do not yet require a fully coherent model with a fixed set of neural areas to support its behavioral predictions. Second, such models, if successful, are a concrete step towards realizing brain-machine-interfaces. For instance, if a model could precisely predict which micro-stimulation patterns lead to which visual percepts, then model-guided stimulation patterns could be used to design visual prostheses to help individuals with lost or impaired vision.

In this study, we ask how well particular ANN models of the ventral stream predict the behavioral effects of direct neural perturbations of IT cortical tissue (Fig. 1). In particular, our contributions are:

1. We develop perturbation modules that formally define the effects of different types of neural perturbations (micro-stimulation, optogenetic and muscimol suppression) on *neural* activity, based on previous biophysical studies.
2. We convert results from four studies measuring the *behavioral* effects of direct IT neural perturbations into a total of nine benchmarks for model evaluation.
3. Combining contributions 1 and 2, we ask how well current topographic ANN models predict the nine empirical benchmarks. We report here that topographic ANN models co-trained with a spatial loss *qualitatively* predict all effects measured in the contra-lateral hemifield. At the same time, these models do not *quantitatively* predict all of those results, and they are unable to explain differences in bilateral processing.

**Fig. 1.**
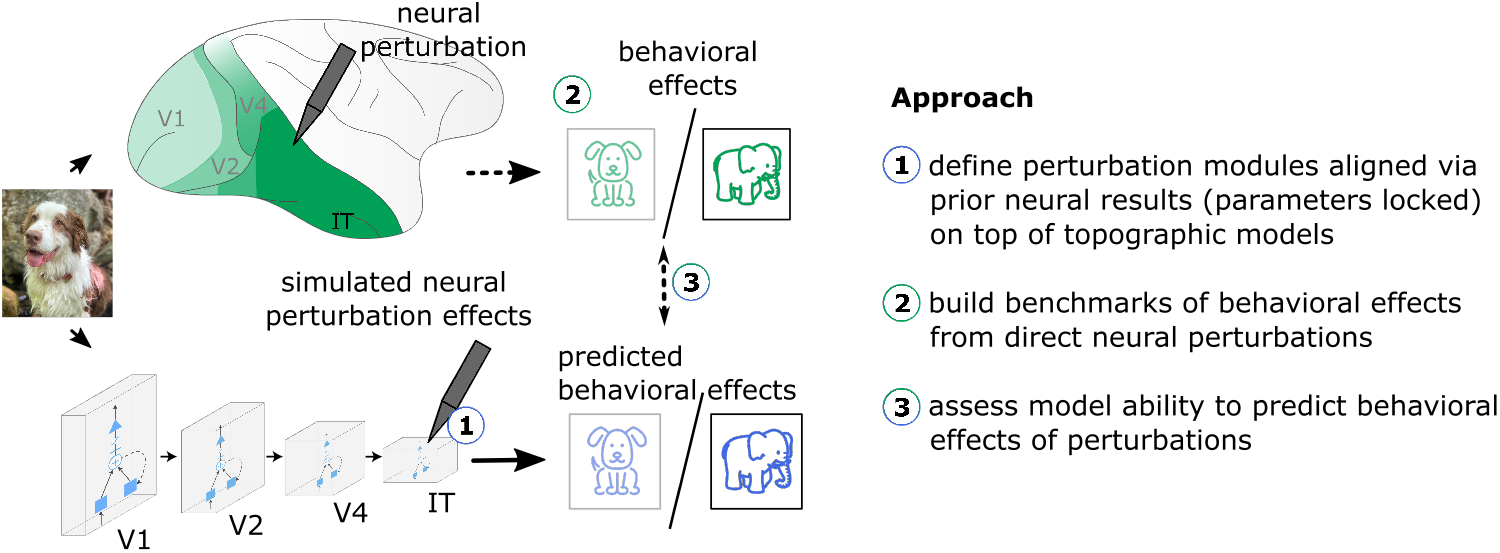
Predicting the behavioral effects of direct neural perturbations with computational models. Direct neural perturbation experiments change neural activity while primate subjects process visual stimuli, and measure the resulting changes in behavior (top, green). To engage with such causal studies with computational models (bottom, blue), we ➀ build fully specified perturbation modules that dictate how neural activity changes as a consequence of neural perturbations, with all parameters locked down by neural recordings, ➁ convert neural perturbation experiments and the observed behavioral effects into benchmarks suitable for model testing, and ➂ test model variants with different strategies of topographically assigning neurons to an artificial tissue sheet by evaluating model predictions of behavioral changes from different neural perturbations against primate experimental observations.

## Results

The rationale of our overall modeling approach is to assume a direct correspondence between the stages of neural processing in artificial neural network (ANN) models and the primate visual ventral stream, with image input, and a downstream behavioral output. The ANN ventral stream models we consider here all have a topography where neurons are organized into 2D model “tissue”, but use different strategies for the spatial assignment. To simulate the effects of direct neural perturbations on the neural elements of a given ventral stream model, we developed a set of perturbation modules (Fig. 1). Each module takes as input perturbation parameters (e.g. the amount of muscimol in an injection, or the current of a micro-stimulation) and outputs the resultant change in neural activity as a spatial profile. That spatial profile can then be applied to a topographic ventral stream model at a particular area of the model (e.g. the layer corresponding to IT) to generate the model-predicted effects of that perturbation. We developed the parameters of each perturbation module based only on the previously-reported neuronal effects of micro-stimulation, optogenetic, and muscimol perturbations. In other words, none of these parameters were tuned to fit any behavioral effects that we later test the models on.

To test these models, we survey published studies that test how neural interventions in primate inferotemporal cortex affect core object recognition behaviors. From these studies we adopt five different experiments based on criteria following replication standards in the field, resulting in nine qualitative benchmarks. After developing the three perturbation modules and locking down all ventral stream model parameters, we then evaluate how well different models predict the experimental observations codified in the nine benchmarks.

### Perturbation modules based on neural data

To engage ventral stream models with tests of direct neural interventions, we first d evelop *f ully s pecified* pe rturbation modules that formally describe the *neural* effects of perturbations, i.e. how neural activity is altered by the application of a particular perturbation at a given location. To determine the parameters of the perturbation modules, we use neural datasets – separate from behavioral datasets that we later test the models on. For all kinds of perturbations, we seek to determine three key parameters: 1. the kind of change to neural activity, e.g. additive / relative to the rate code activity in model units (∼ corresponding to spike rates); 2. the magnitude of neural activity change at the center of the perturbation; and 3. the spatial profile of that change.

#### Muscimol suppression

Arikan et al. (2002) characterized both magnitude and spread of neural suppression following muscimol injection. At the center of injection, neural activity is completely suppressed. With increasing distance from the injection site, activity suppression decreases. We translate the qualitative judgements into a quantitative Gaussian spread of suppression (𝒩 (*μ* = 1, *σ* = 1.17*mm*), Fig. 2c), taking “Complete Suppression” to mean 100%, “More” 67%, “Less” 33%, and “None” 0%.

**Fig. 2.**
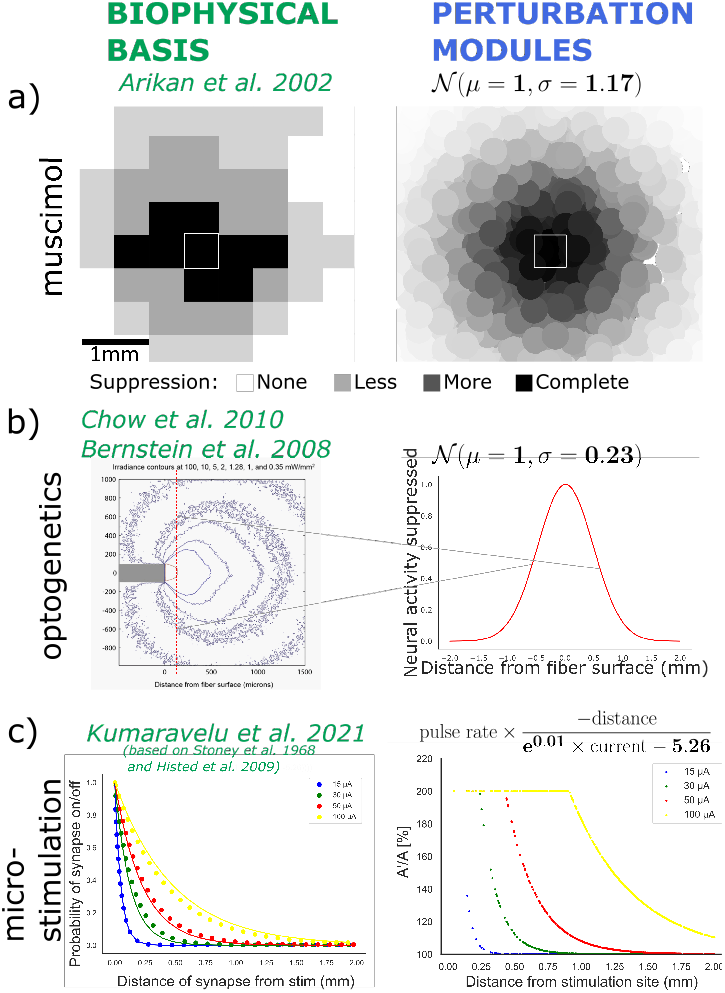
Perturbation modules aligned via prior neural results. We develop perturbation modules that fully specify how direct neural perturbations affect model neural activity. Each module’s parameters are locked down by prior neural experiments that measure how neural recordings change as a function of perturbation parameters. **Muscimol suppression, a)** We use the neural suppression effects observed by Arikan et al. (2002) to fit a G aussian f all-off t o t he scale-dependent relative suppression. **Optogenetic suppression, b)** Chow et al. (2010) observed a Gaussian spread of light irradiance under different applications of mW/mm^2^, and Bernstein et al. (2008) found that “a radiant flux of 1 m W/mm^2^ will activate approximately 50% of molecules”. We combine these two findings into a Gaussian fall-off of relative suppression, and directly translate activation of molecules into suppression of neural activity. We set the magnitude to 1 for all output powers *≥* 6 mW/mm^2^ following an observation that 100% of spikes are silenced in this setting (Chow et al., 2010). **Micro-stimulation, c)** We adapt the neural effects of micro-stimulation to biophysical modeling by Kumaravelu et al. (2021) who unified experimental observations (Stoney et al., 1968; Histed et al., 2009). In our perturbation module, the magnitude of added neural activity is determined by the pulse rate (Hz), and the spatial spread follows an exponential fall-off parameterized by the current (mA).

#### Optogenetic suppression

For optogenetic neural perturbation, biological neurons are prepared for a direct perturbation by first infecting them with a virus that carries the genetic material to express a particular opsin protein. The perturbation is later executed by activating the opsins with an optical laser pulse, which then either suppresses or enhances (depending on the opsin) neural activity in infected neurons. Chow et al. (2010) measured light irradiance distances from the fiber tip for different strengths of light (*mW/mm*^2^). Bernstein et al. (2008) further connected the radiant flux to molecule activation, observing that 1*mW/mm*^2^ activated ≈50% of molecules. We translate this into a Gaussian spread of molecules activated as a function of distance from the fiber tip, fit to the irradiance contours (𝒩 (*μ* = 0.5, *σ* = 0.23), Fig. 2b). In this study, we focus only on optogenetic suppression and assume that the neural effects of Arch opsins are comparable (Arch-GFP in Chow et al. (2010), ArchT in Afraz et al. (2015)). In terms of magnitude, Chow et al. (2010) found that for 6*mW/mm*^2^ irradiance, 100% of spikes are silenced. We thus assume that for ≥ 6*mW/mm*^2^ irradiance, the magnitude of silencing is full neural suppression. From the fiber tip, neural activity is then scaled relatively according to the fall-off of molecule activation.

#### Micro-stimulation

Electrodes (single shank as well as multishank arrays) can deliver electrical pulses that activate local clusters of neurons. To compute the effect of microstimulation on neural activity, we follow the biophysical model developed by Kumaravelu et al. (2021) which unifies effects observed experimentally by (Stoney et al., 1968) and (Histed et al., 2009). The two primary stimulation parameters we consider are the pulse rate (Hz) and current (mA): The pulse rate specifies the relative increase of neural activity magnitude, while increased current increases the spatial spread of activity increase. More specifically, the relative activity increase for a neuron a certain distance (mm) away from the stimulation site is governed by an exponential fall-off (Fig. 2a), and clipped such that each unit activity is at most doubled (Ponce et al., 2019).

### Spatial assignment of model IT neurons

In this study, we consider the spatial profile of perturbations in the immediate vicinity around the stimulation or injection site. To simulate this in models of the ventral stream, model neurons must first be assigned to *x* and *y* locations within artificial cortical tissue (e.g. a simulated IT cortical tissue sheet). This simulated tissue sheet is the basis for simulating how neural activity in the ventral stream model is affected by the various perturbations. We consider three ways of building a model tissue sheet. In each case, we assume the underlying ventral stream model has already specified its cortical areas (e.g. it has an IT area containing a set of simulated IT neurons). Specifically, following Lee et al. (2020), we use the same AlexNet base architecture to generate all three TDANN model versions. Note that, for the first two procedures below, the functional roles of all simulated neurons are already determined by pre-training (see Methods), prior to the spatial tissue mapping.

### TDANN via random arrangement

Here, we assign each of the neurons in a given cortical area to random locations within the artificial tissue sheet. This does not change the functional role of any simulated neuron.

### TDANN via swapopt

This procedure optimizes the relative locations of all neurons within the artificial tissue sheet according to a spatial loss Margalit et al. (2020), starting from initial random locations. This again does not change the functional role of any simulated neuron.

### TDANN via spatial co-training

Neurons are assigned to fixed (random) locations, and the parameters of the model are optimized jointly with a classification and a spatial correlation loss (Lee et al., 2020). This is the only model type in which the functional roles of the neurons are impacted by the spatial arrangement.

#### Quantifying model tissue to primate tissue spatial and neural alignment

As a basic test of the *spatial and neural* alignment of a model’s tissue to biological IT tissue, we used a known property of IT tissue: spatially-nearby IT neurons tend to have similar functional responses and spatially separate neurons have different response profiles. We used the same strategy of prior work and compute the functional similarity of pairs of IT neurons as the Pearson correlation over responses to a common set of images, and then plot that correlation as a function of the spatial distance of the two neurons. The same procedure was applied to each IT tissue model and then compared with the same empirical measure from primate IT previously reported Lee et al. (2020). The image set that was used to measure functional similarity consisted of 5,760 naturalistic images from Majaj et al. (2015) along with the associated IT neural recordings, though unpublished results suggest that the particular images set is not critical to this measure. For models, we simulate sub-sampling from Utah array placements similar to the primate recordings by sampling model IT neural activity from 3.6 × 3.6*mm* randomly placed “electrode” grids.

As expected, models with random spatial organization exhibit no distance-dependent change in response correlations. In contrast, models with spatial co-training show a qualitatively similar trend to the primate data: neurons that are close together respond similarly, whereas neurons that are far apart respond dis-similarly.

To quantify the spatio-functional alignment of model to primate IT, we use a Kolmogorov-Smirnov (KS) metric: for each distance bin of 0.8*mm*, we aggregate response-correlation values within that bin. The KS similarity then evaluates how likely samples are to be drawn from the same underlying distribution. The final score is the median over bins. Despite a qualitative match, TDANN models with spatial co-training do not perfectly match the IT data (e.g. sample model in Fig. 3c, ceiling-normalized score 0.75 with a raw KS similarity of 0.58 ± 0.06 relative to a monkey-to-monkey ceiling of 0.77± 0.11), suggesting that the TDANN model still differs from how primate IT tissue is organized.

**Fig. 3.**
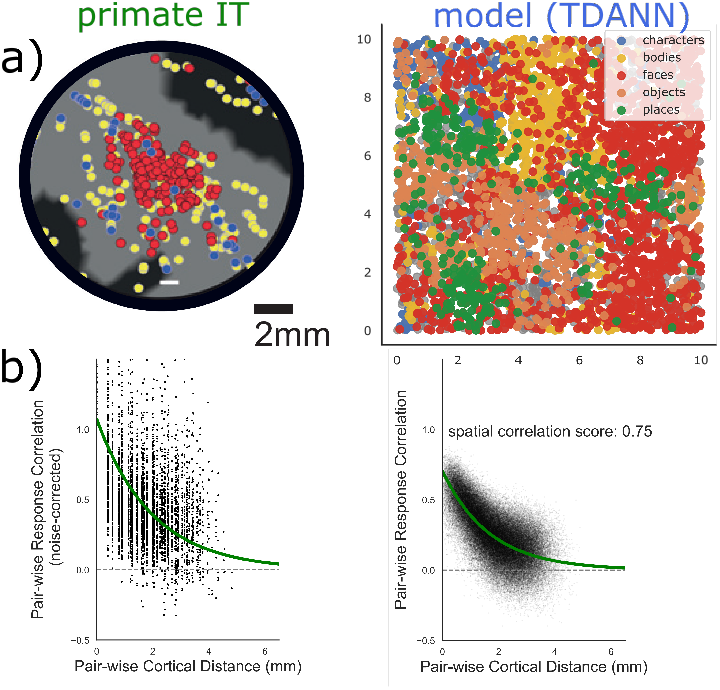
Building a model tissue sheet. Comparison of neural activity in primate IT (left, Majaj et al. (2015)) and the “IT” layer of a topographic model (right, “TDANN via spatial co-training” (Lee et al., 2020)): **a)** Neurons in primate IT and in TDANN models both demonstrate spatial clustering when assessed for category preference. Left panel shows results from a neurophysiology study of IT cortex showing the tissue locations of neural sites that responded more to faces than non-face objects (red), neural sites that respond more to non-face objects (blue), or had little response preference (yellow). Adapted from Lee et al. (2020). Right panel shows the tissue map of the AIT layer of a single TDANN model, where the color indicates the prefered category among a set of tested categories. Note that spatial scales are comparable between primate and model IT, for example face-preferring clusters are 2-4 mm in diameter. **b)** Lee et al. (2020) utilizes a decaying exponential relationship to quantitatively describe the pair-wise response correlation of primate IT sites and their cortical distance – i.e., nearby sites are more similar to one another, and distant sites are dis-similar. Left panel shows the pairwise correlation of sites over an image set from Majaj et al. (2015, y-axis). Right panel shows the corresponding effect in a TDANN model.

**Fig. 4.**
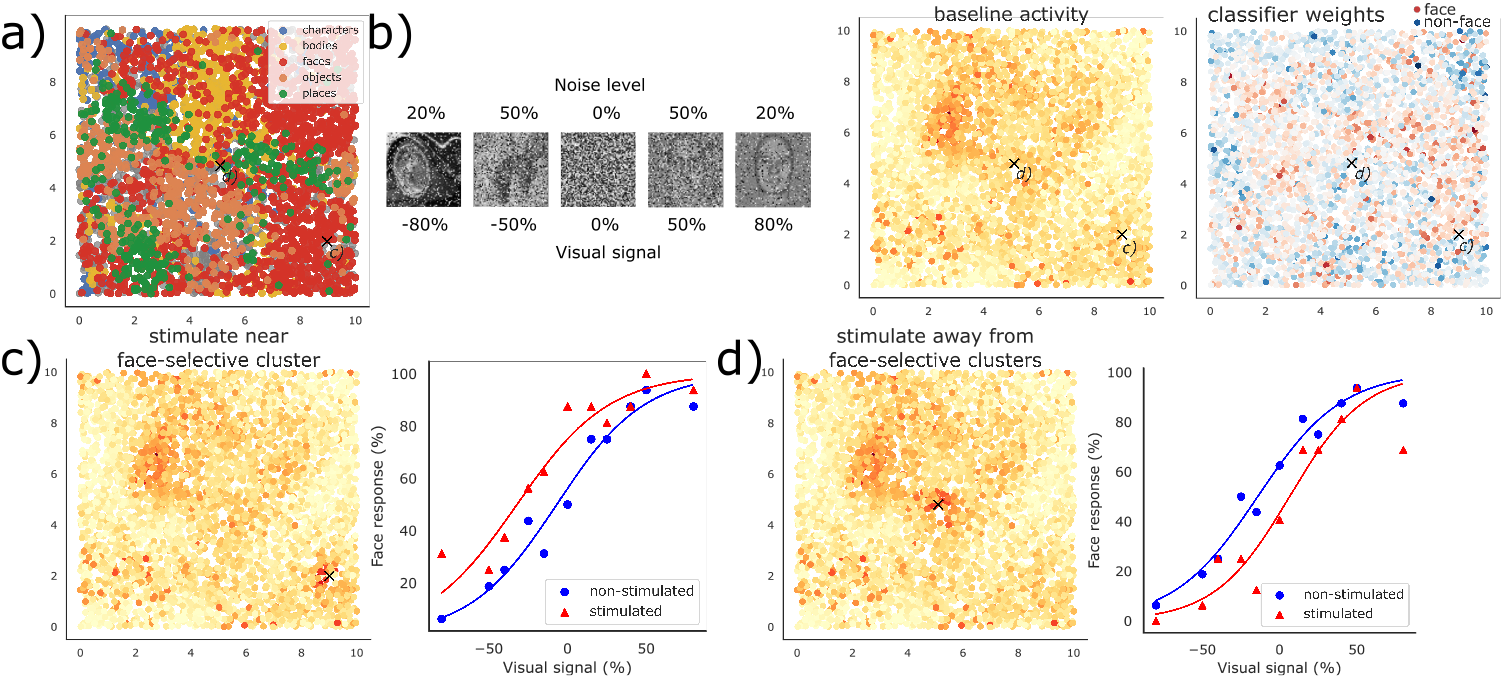
Example micro-stimulation perturbation experiment on an individual TDANN model. **a)** AIT tissue layer from a TDANN model (determined via spatial co-training; image reproduced from cf. Fig. 3). The X at the center and the X at the lower right indicate the tissue locations of two simulated micro-stimulation experiments (c and d). **b)** Left: Stimuli used in the non-human primate micro-stimulation experiments of Afraz et al. (2006) In each row, the stimuli gradually transition from non-face object (left) to face (right) and this was defined by the experimenters as a face “visual signal” axis which ranges from -100% (minimal face signal) to +100% (maximal face signal). Face images are derived from computer-generated faces (Moeller et al., 2017) for visualization purposes. Middle: The same stimuli were presented as visual inputs to the TDANN ventral stream model and here we plot the average activity over all stimuli. Lighter and darker colors respectively indicate less and more active unit activity. Right: Simulating the behavioral training of the non-human primates, a linear classifier was trained to report face vs. non-faces across 510 separate images (based only on the output of the TDANN model AIT layer; see Methods). After decoder training, weights are frozen for the microstimulation simulations. To illustrate how the TDANN model AIT supports the final behavioral decision (face vs. non-face) we plot the classifier weights (the more red, the more a positive activation of that unit contributes to the behavioral decision to report face). **c)** Example of a simulated micro-stimulation direct IT neural perturbation experiment. In this example, the electrical stimulation site was near a face-selective cluster of simulated neurons (i.e. inside a cluster of red units in panel *a*, and with mostly positive face classifier readouts in panel *b*). Left: The simulated effect of the micro-stimulation (generated via the micro-stimulation perturbation module, Fig. 2; see text) results in an increase in activity of simulated neurons near the site (cf. with panel *b* middle). Right: The behavioral psychometric curve predicted by this model for both the non-perturbed and the perturbed (i.e. micro-stimulated) conditions (see Methods for details on how this curve was generated). This TDANN model predicts that, at this particular micro-stimulation site, there will be a shift in the behavioral psychometric curve to the left (of the magnitude shown from blue to red). This shift means that, with micro-stimulation, the model is more likely to report “seeing” a face for any given level of face signal. **d)** Example of a different micro-stimulation site for the same TDANN model. In this case, the site is not near a strong face cluster (see *a, b*). In this case, the model predicts a shift of the psychometric curve to the right. All plotting conventions are the same as in panel *c*.

### Building causal intervention benchmarks

With the models (topography + perturbation modules) locked down, we next tested how well they could predict experimental data “out of the box.” Based on previously published studies, we built perturbation benchmarks from each paper’s central claims and supporting experimental data. To select which studies and claims to adopt into benchmarks, we first surveyed the literature, obtained experimental stimuli, and then adopted each paper’s central claims.

#### Literature survey

We here focused on the effects of perturbations in inferotemporal cortex (IT) on foveal (central 8 degrees) core object recognition behaviors. This lead us to exclude perturbations in early visual cortex (e.g. Jazayeri et al., 2012), learning effects (e.g. Kawasaki and Sheinberg, 2008)), as well as large stimulus presentations (e.g. Sadagopan et al., 2017). For most of the remaining studies, we were able to obtain experimental stimuli, leaving us with four core studies: Afraz et al. (2006, 2015); Moeller et al. (2017); Rajalingham and DiCarlo (2019).

#### Adoption criteria

To select which observations to convert into quantitative benchmarks, we followed two key criteria: First, we focus on those experimental data that support each study’s central claims (i.e. those highlighted in the abstract). We do this under the assumption that a publication’s core claims are the ones that are the best vetted and most likely to be true. Second, we use the field’s standard criteria of replication in at least 2 monkeys, i.e. we excluded any claims and data that were tested in only 1 monkey. In total, we adopted 9 claims and supporting data from 5 different experiments into quantitative benchmarks (Fig. 5 and Table 1).

**Table 1.**
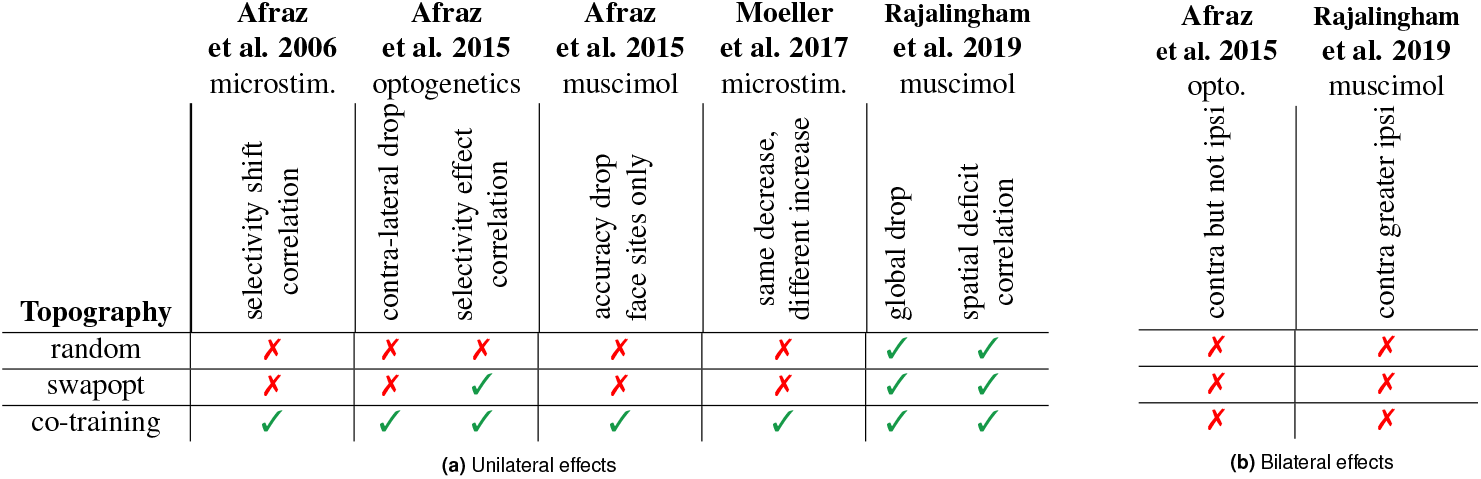
Benchmarking topographic model variations. **a)** We score topographic models (with different strategies for building an artificial tissue sheet) on the 9 perturbation benchmarks. Random arrangement of units matches the un-changed ipsi-lateral effect in Afraz2015 as well as two benchmarks in Rajalingham2019, but fails all others (2/7). Arranging neuron locations after standard training (swapopt, Margalit et al., 2020) improves slightly over random topography (3/7 ✓). The TDANN with spatial co-training (Lee et al., 2020) qualitatively predicts all unilateral effects (7/7 ✓, see Fig. 5 for details). **b)** Since none of the models feature separate hemispheres, all fail to predict bilateral processing where perturbations lead to more behavioral changes with contra-lateral compared to ipsi-lateral images.

**Fig. 5.**
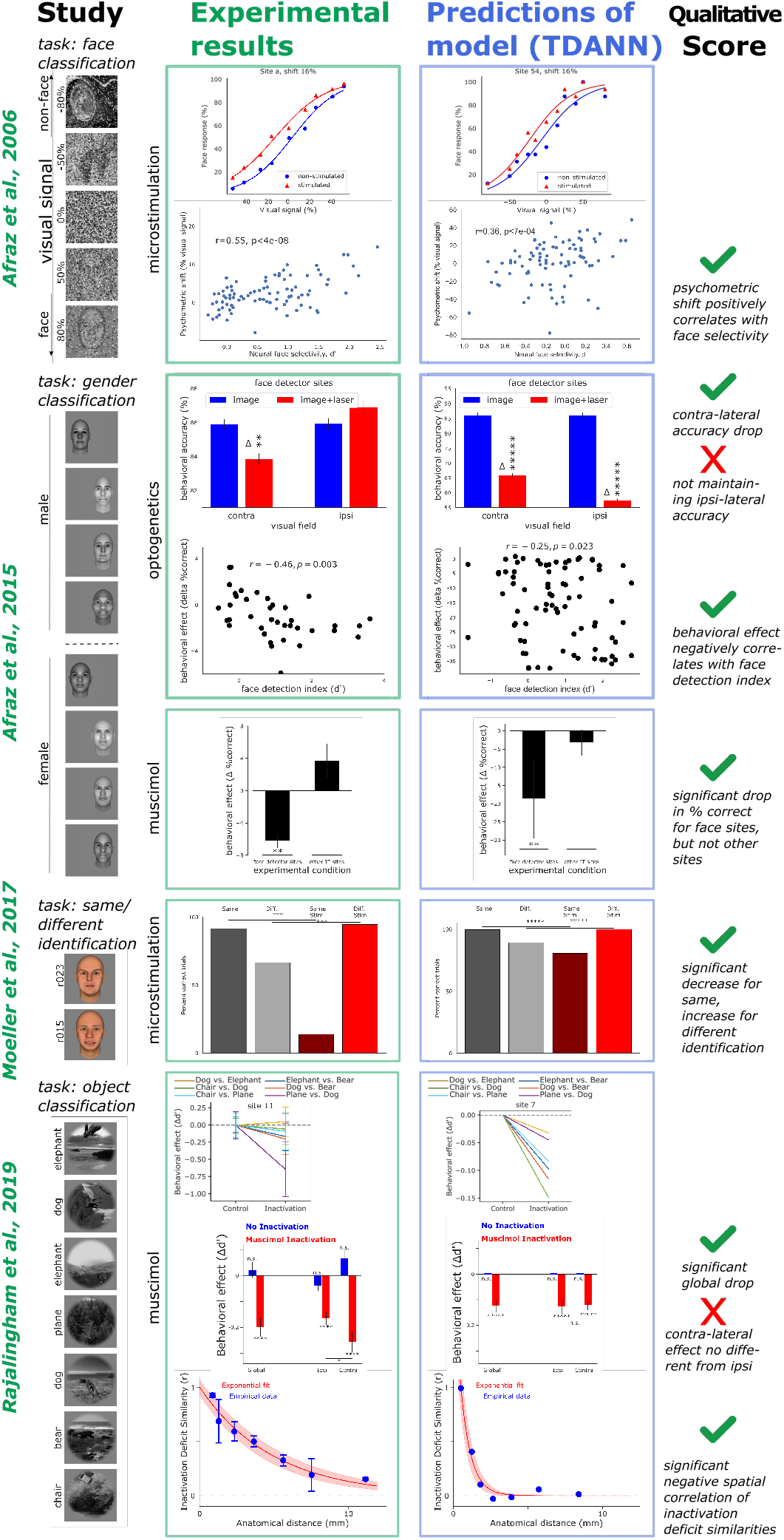
Benchmarking a topographic model on its ability to predict the behavioral effects of neural perturbation experiments. For each benchmark, the leftmost column visualizes the experimental paradigm, including task and stimuli (all visualized faces are computer generated). Experimental results (green boxes) display primate measurements in support of the respective claims. Example model predictions (blue boxes, TDANN via spatial co-training), are derived from testing the model in the same way as the primate experiments (Methods). The final “Score” column denotes whether model predictions qualitatively match experimental results. **Afraz2006:** Subjects were presented with images ranging from faces, to pure noise, to non-faces, and had to categorize stimuli as face or non-face. *Top row* : Sample shift in the psychometric curve of face responses with stimulation (red). *Bottom row* : Dependence of the psychometric shift on site’s face selectivities. **Afraz2015, optogenetics:** Subjects had to classify face images as either ‘male’ or ‘female’. *Top row* : Overall change in behavioral accuracy with no optogenetic suppression (blue) and for contra- and ipsi-lateral presentation of stimuli. *Bottom row* : Dependence of the per-site behavioral effect on the site’s face selectivities. **Afraz2015, muscimol:** Same task as for the optogenetics experiments, but neurons are suppressed with muscimol injection instead of optogenetics. The plot shows the change in behavioral accuracy when suppressing face-selective sites (*d′ >* 1) or other sites. **Moeller2017:** Subjects were presented with two face images in succession and had to classify whether they were the same or different identity. Gray bars correspond to the behavioral accuracy in same/different trials without stimulation, red bars with stimulation. **Rajalingham2019:** Subjects were presented with objects from 6 categories in random viewing conditions and had to classify the resulting gray-scale images in match-to-sample tasks. *Top row* : Behavioral effects in 6 sub-tasks (e.g. “Dog vs. Elephant”) when injecting muscimol in a sample site. *Middle row* : Summary of behavioral effects for all images (“Global”), and ipsilateral/contralateral images (“Ipsi”/”Contra”). *Bottom row* : Similarity in behavioral deficits of site pairs as a function of the anatomical distance between injection sites.

### A topographically organized model qualitatively predicts behavioral perturbation effects

We now apply the perturbation modules to the topographic models so that the overall model (Fig. 1): processes visual input through a hierarchy of feature transformations, perturbs neural activity in model IT according to a particular direct neural intervention based on the perturbation modules (➀, Fig. 2) and its spatial assignment of neurons (➁, Fig. 3), and finally uses the resulting IT activity to make a behavioral response in a given task. The behavioral decoder is a simple linear readout, taking as input the (perturbed) IT features and producing class choices. This decoder is fit for each benchmark separately (see Methods).

#### Simulating the effects of direct neural perturbations on behavior

Fig. 4 shows sample effects of micro-stimulation on the topographic TDANN model. Under the *Afraz2006* experimental paradigm, images of faces and non-faces are shown, each distorted with a random noise level between 20-100%. Non-face stimuli with little noise (negative visual signal) are consistently classified as ‘non-face’ while face stimuli with little noise (positive visual signal) are correctly classified as ‘face’ (Fig. 4c and d, blue lines).

In the *Afraz2006* experiment, micro-stimulation is then applied to different sites across IT. Stimulating model sites leads to an increase of neural activity surrounding the site, in addition to the image-elicited activity (Fig. 4c and d left, sample site). Just like with primate experiments (Afraz et al., 2006), this increase in firing rate shifts the psychometric curve of face responses such that subjects are now more likely to respond ‘face’ (Fig. 4c, red line).

Since the models we test here are imageas well as perturbation-computable, they can applied to a variety of different experiments. Depending on the stimuli and perturbations as well as the topographic model, these experiments will change model neural activity in different ways, resulting in different behavioral predictions. We next test whether these model-predicted behavioral changes under different experimental paradigms match the corresponding primate observations. We primarily focus on the predictions by the topographic model with spatial co-training, and present all model results in Table 1. Model ability to predict experimental results is summarized in a qualitative score: ✓ denotes qualitative alignment, ✗ denotes mis-alignment.

#### Shifting psychometric curves for face classification with micro-stimulation (Afraz et al., 2006

For stimuli ranging from non-faces (visual signal*<* 0) to noise (0% visual signal) to faces (visual signal*>* 0), Afraz et al. (2006) measured the psychometric curve of face responses without and with micro-stimulation. Micro-stimulation shifts the psychometric curve to the left, making primate subjects more likely to respond ‘face’ to the same visual stimuli (Fig. 5 top row, green box). Across stimulation sites with different face selectivities, the shift of the psychometric curve is more pronounced towards faces when a site is more face-selective (*r* = 0.55, *p <* 0.00001).

When stimulating sites in the model with random topography, the psychometric curve barely shifts. The shift of the psychometric curve further does not depend on the face selectivity of stimulated sites (*rn*.*s*., Table 1). Intuitively, when sites are not clustered by functional preferences, stimulation results in non-specific effects: if a neuron is face-selective, neighboring neurons might not be selective to faces, such that the behavioral read-out is more scattered around tissue. Stimulating at face-selective sites might thus only affect a small portion of face-selective neurons, resulting in a small and non-specific shift in the psychometric curve.

Stimulating the topographic model built with spatial co-training on the other hand leads to bigger shifts in the psychometric curve, similar to those observed experimentally (Fig. 5, blue box). When neurons are clustered according to their functional preferences, stimulating a face-selective site here has a high chance of stimulating other face-selective neurons in the vicinity, leading to larger and more selective shifts in the face responses. The extent of the psychometric shift depends on the face selectivity of the stimulated site, just like in the primate experiments, although the correlation is smaller (*r* = 0.36, *p <* 0.001), and the psychometric shifts are larger (up to∼ 40% visual signal in the model compared to *<* 20 in the primate experiment).

#### Impairing face gender classification with optogenetic suppression (Afraz et al., 2015, optogenetics)

In Afraz et al. (2015), the authors tested the causal involvement of primate IT in the discrimination between faces. Specifically, subjects are presented with a male or female face and asked to report the perceived gender.

When neural activity was optogenetically suppressed, the behavioral accuracy decreased from 86% to 84% in the contralateral visual field (*p <* 0.0001). Suppressing at more face-selective sites resulted in a more pronounced drop in behavioral accuracy (*r* = − 0.46, *p <* 0.003).

The topographic model with spatial co-training reproduces the drop in behavioral performance, although its baseline performance as well as the drop are higher (92% to 88%, *p <* 0.0001). The model also predicts the dependence of the behavioral effect on the face selectivity of the optogenetic suppression site (*r* = − 0.25, *p <* 0.05). Model-predicted behavioral effects were generally larger than observed in the primate experiments.

#### Impairing face gender classification in face detector sites with muscimol suppression (Afraz et al., 2015, muscimol)

The same study tested the behavioral effects of neural suppression with muscimol.

This experiment revealed a dependence of reduced face gender classification performance on sites’ face selectivities: suppressing face detector sites lead to a greater behavioral effect (-5%, *p <* 0.01) whereas suppression of other IT sites (face selectivity *d′* ≤ 1) lead to no significant decrease in behavioral performance.

When model sites are suppressed with muscimol, the model correctly predicts a drop in accuracy for face detector sites (*p <* 0.01) with no significant drop for other IT sites. Again, the predicted drop is larger than the experimentally observed behavioral changes. Contrary to the model, the primate data also showed a slight improvement of behavioral accuracy when other IT sites were suppressed, but this effect was non-significant.

#### Biasing face identification with micro-stimulation (Moeller et al., 2017)

Moeller et al. (2017) tested the effects of micro-stimulation on the perception of faces. Subjects were presented with two images consecutively, and reported whether the two images were the same or different identity. The central claim of the paper tests the recognition of same/different face identities, without and with stimulation.

When micro-stimulating inside an IT face patch, primate ability to detect ‘same’ face identities strongly decreased (*p <* 0.001), whereas the detection of ‘different’ identities became more accurate (*p <* 0.001). Overall, primate performance thus still decreased, but with a differential effect between ‘same’ and ‘different’ trials.

When the topographic model with co-training is stimulated in an IT face patch (see Methods) during face-identity trials, it correctly predicts the decrease in ‘same’ identity (*p <* 0.00001) and the increase in ‘different’ identity detection (*p <* 0.00001), although with smaller effect sizes.

#### Deficits in object categorization with muscimol suppression (Rajalingham and DiCarlo, 2019)

Rajalingham and DiCarlo (2019) tested the effects of reversible muscimol suppression on core object recognition performance in match-to-sample tasks. Subjects were presented with a test image, followed by a 2AFC screen with another image of the same category as the test image and a distractor image. The combination of test and distractor categories made for 10 sub-tasks such as “elephant vs. bear”, “dog vs. bear”, etc.

Muscimol suppression at each site selectively affected primate performance on sub-tasks, i.e. some sub-tasks were affected more than others. On average, behavioral performance decreased by Δ*d*′ = − 0.2 (*p <* 0.0001), notably with a significant difference between ipsi- and contra-lateral object images. Correlating deficit patterns (over sub-tasks) across sites, the study further determined that inactivating nearby sites lead to similar deficits whereas inactivation of distant sites showed dis-similar effects which was fit with an exponential model (Spearman *r* = 0.36, *p <* 0.001).

While the topographic model with spatial co-training exhibits a selective effect of muscimol suppression on the different sub-tasks, the sparsity of affected sub-tasks on average is lower. The model predicts the overall behavioral deficit well (*p <* 0.00001), but, unsurprisingly, does not exhibit any difference between ipsi- and contra-lateral object images. Finally, the model correctly predicts the dependence of inactivation deficit similarity between sub-tasks on the anatomical distance of inactivation site pairs, although the similarity seems to fall off more quickly than in the primate experiments.

#### Topographic model with spatial co-training predicts all unilateral qualitative effects

In summary, the topographic model with spatial co-training qualitatively predicts 7 out of 9 experimental observations, in different task settings (ranging from face to object identification and categorization) as well as different perturbation methods (micro-stimulation, optogenetic, and muscimol suppression). The 2 benchmarks that this model fails to predict both correspond to differences in contra- and ipsi-lateral effects. Despite the qualitative successes, quantitative predictions are often questionable which we later investigate in more detail (Fig. 6).

**Fig. 6.**
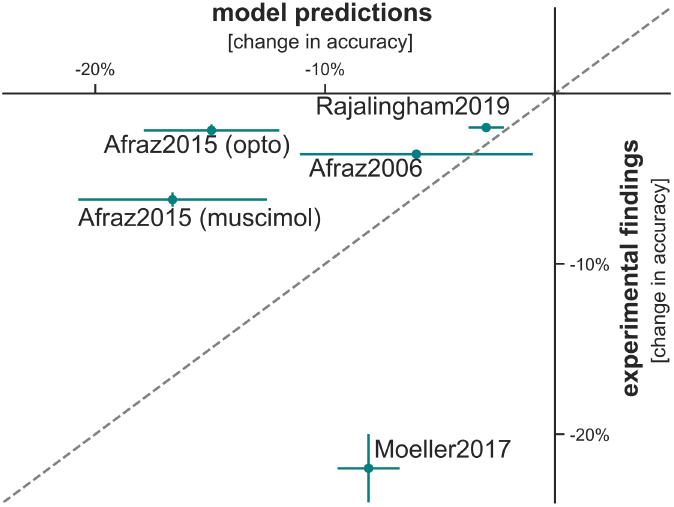
Quantitative misalignment. While the TDANN via spatial co-training predicts all unilateral qualitative effects (Table 1a), it fails to capture exact quantitative outcomes. We measure the overall behavioral effect as the difference in balanced accuracy between no perturbation and perturbation conditions. Note that we report the overall difference in task performance, i.e. for *Moeller2017* we combine same and different trials. All primate and model experiments observe a detrimental (negative) effect on balanced accuracy. Error bars are s.e.m. over subjects (5 TDANN instances from different seeds for the model, 2 macaque monkeys for biological experiments).

### Benchmarking topographic model variation

We next evaluated model variants with different topographic strategies for the artificial tissue sheet of neurons in IT. Compared to the main models used in this study which co-train with a categorical and a spatial loss, a model trained with a categorical loss only and random arrangement of units in x and y locations fails to predict most perturbation benchmarks (2/9 ✓). The post-hoc assignment of neurons to spatial locations according to a spatial correlation loss (‘swapopt’ Margalit et al., 2020) similarly lead to poor response profiles (score 0.18, cf. Fig. 3) and predicted slightly less than half of the perturbation benchmarks (3/9 ✓).

### Quantitative misalignment

Going beyond the central *qualitative* claims typically put forth in experimental papers, we sought to evaluate the quantitative alignment of model predictions to experimental findings. Specifically, we used the best-performing model (“TDANN via spatial co-training”) and evaluated the magnitude of its predictions against the magnitude of behavioral effects observed in prior studies. This evaluation builds on the assumption that quantitative findings are robust across different experimental parameters, such as the exact location of neural perturbations. Most studies considered here estimate the robustness of effect magnitudes and report error bars across subjects, making us confident that we could evaluate the overall trends of model predictions versus biological findings.

While model predictions are qualitatively aligned to all unilateral effects we evaluate here, the quantitative predictions are consistently misaligned. Of the five experiments with unilateral effects, the models (here from 5 different seeds) over-predict the average change in accuracy for three of them: For psychometric shifts in face classification following microstimulation (Afraz et al., 2006), the models predict an overall shift of − 6 ± 5% while the authors found a − 4 0. ± 2% shift in their experiment (error estimates are standard error of the mean with *n* = 2 primate and *n* = 5 model subjects). For a deficit in behavioral accuracy following contralateral optogenetic suppression of face detector sites, the models predict a delta of − 15 ± 3% but experimentally only a − 2 0. ± 3% deficit was measured. Similarly, the models predicts a − 17 ± 4% accuracy deficit with muscimol suppression of face detector sites but experimental data suggests only a tal error estimates were only available for *d* ′ measures). Only − 6 0. ± 4% difference (Afraz et al., 2015). In an object classification task, muscimol suppression of individual sites (Rajalingham and DiCarlo, 2019) leads to a model prediction of an overall −3 ± 0.8% deficit in balanced accuracy which is similar to the experimental observation of −2% (experimen-for one experiment where sites in a face identification task are microstimulated (Moeller et al., 2017) do the models underpredict how strongly primate behavior is affected: the models predict a − 8 ± 1% deficit in identifying same faces while the experimental data points to a − 22 ± 2% deficit. Note that error bars especially across model subjects are often large, leaving the exact quantitative discrepancies unclear. Overall, the quantitative predictions of behavioral changes in accuracy seem moderate at best.

## Discussion

This work enables artificial neural network models of primate visual processing to engage with causal perturbation experiments. Making perturbation studies accessible for model evaluation, we convert the nine central claims in five different primate IT perturbation experiments into model benchmarks. Based on neural recordings, we developed fully specified perturbation modules that translate micro-stimulation, optogenetic, as well as muscimol suppression into changes in neural activity. Applying these perturbation modules to topographic model variants, we found that an ANN co-trained with a spatial loss predicts all unilateral benchmarks, but cannot engage with bilateral benchmarks and is quantitatively misaligned.

### Model successes: qualitative predictions of all unilateral experimental claims

Across four different studies, the model correctly predicts all qualitative unilateral effects. Note that model parameters were locked down entirely based on task and topographic optimization (for the underlying ANN) and separate experiments measuring the effects of perturbations on neural activity (for the perturbation modules). All of these behavioral effects are thus novel predictions by the model that, without any fitting, align with biological data.

### Model incompleteness: bilateral hemispheric effects

Despite the assignment of model units to spatial tissue, the topographic models tested here optimize for a single “hemisphere”. In other words, there is no bilateral processing in these models. As a consequence, the models make the same predictions for contra- and ipsi-lateral images. This is different from primate studies where behavior is typically affected more by contra-lateral perturbations.

### Model failures: quantitative mis-predictions

While the studies we focused on here did not emphasize quantitative effects as their main claims, we believe effect magnitudes are an important component and compared the deltas in behavioral performances between primate and model experiments. The best TDANN type (spatial co-training) is not aligned to quantitative results and over-as well as under-predicts changes in behavioral performance (Fig. 6). Several modeling choices could be the cause of this mismatch:

#### Perturbation modules

While we based the perturbation modules on prior neural observations (Fig. 2), we still had to make some assumptions: for micro-stimulation, we assumed that a higher pulse rate would lead to a higher increase in firing rate with a linear relationship. However, we could not find experimental data that established the precise relationship between pulse rate and firing rate. With optogenetic suppression, we had to assume several links from laser output power to light irradiance to molecule expression to neural firing rates. While we used experimental findings to create those links, the causal chain from optogenetic activation to spike rates is not directly tested by these experiments. The effects of muscimol suppression on neural activity were the most well captured experimentally, with a clear link from virus injection to neural suppression levels.

#### Topography

The core free parameters of the model variants are in their topography: training the model to learn spatially dependent representations showed a clear improvement over random assignment of neurons to tissue locations. But even with purely neural measurements, TDANN model tissue is not perfectly aligned with primate IT recordings (Fig. 3). Further improving the artificial tissue sheet as well as functional responses (Schrimpf et al., 2020) could help close the quantitative gap between model predictions and primate experiments.

#### Fixed decoder

Finally, we here built end-to-end models of representational shifts under an assumption of a fixed decoder. In the behavioral paradigm for the primate experiment, subjects always received rewards for correct trials. Continuing to finetune decoder parameters during the perturbation phase would reduce the behavioral deficit predicted by the model. This would alleviate quantitative mismatches with 4 of the benchmarks, but would make the *Moeller2017* experiment even more of an outlier.

### Brain-machine interfaces

The results in this work make us cautiously optimistic that the computational models we develop are promising to, one day, help with clinical applications. In particular, if we were to discover a model that could precisely predict the behavioral effects of micro-stimulation, this model might be useful for visual prosthetics. Current approaches stimulate in V1 to elicit visual percepts of simple shapes (Beauchamp et al., 2020; Chen et al., 2020; Fernández et al., 2021) – but even with dozens to hundreds of implanted electrodes, eliciting high-level visual percepts from V1 stimulation alone seems to be out of reach for now. Stimulating high-level visual cortex IT (or a combination of e.g. V1 and IT) on the other hand might be more viable for object percepts since the stimulation of few sites might be sufficient for object representations compared to stimulating the entire object outline. Future closed-loop model-guided visual stimulation experiments will hopefully help to determine the feasibility and requirements of high-level visual prosthetics.

The approach, benchmarks, and models presented here lay the foundation for a research path where models of the primate visual ventral stream are routinely tested on an integrative, growing set of qualitative and quantitative benchmarks from a multitude of experimental perturbation studies – which in turn guide and constrain the development of the next generation of causally predictive models.

## ACKNOWLEDGEMENTS

We thank Tiago Marques for helping with the KS-metric, and Binxu Wang and Carlos Ponce for helping with neurons’ maximum firing rates. This work was supported by a Takeda Fellowship (M.S.), a Massachusetts Institute of Technology Shoemaker Fellowship (M.S.), a Friends of the McGovern Fellowship (M.S.), the SRC Semiconductor Research Corporation (M.S., J.J.D.), the Office of Naval Research Grant MURI-114407, and the Simons Foundation Grant SCGB-542965 (J.J.D.).

## Methods

### Benchmarks

All benchmarks attempt to codify the experimental paradigm that published data was collected with. We apply this paradigm to candidate models to assess whether the resulting model predictions align with the corresponding primate observations.

#### Stimuli

Whenever possible, we use the same stimuli to test models that were used in primate experiments. Specifically, for *Afraz2006*, the original images were no longer available, but we re-created (Fig. 5) similar stimuli by sampling 30 face images from Labeled Faces in the Wild (LFW, Huang et al., 2007) and 60 non-face images from ImageNet 2012 (Deng et al., 2009). For each image, we down-sampled to 64 × 64 pixels, converted to grayscale, and added uniform noise for the varying signal levels. For *Afraz2015*, we had access to the same image bank and sampled 400 images for testing (200 for muscimol experiments following the paper). For *Moeller2017* and *Rajalingham2019*, we used the exact same face images as in the original study.

#### Training paradigm

To provide model candidates with training stimuli for the task, we follow the experimental setup as specified in the respective paper whenever possible. Specifically, for *Afraz2006*, we select 510 images of the same distribution for training. For *Afraz2015*, we sampled 400 images of the image bank for training. For *Moeller2017*, we use the images from the experiment, and for *Rajalingham2019*, we use training images from a similar image distribution created by Majaj et al. (2015). Approximating primate juice rewards, labels are provided directly with each stimulus.

#### Perturbation sites

We again attempt to perturb in the same way as was specified in each paper’s methods. Specifically, for *Afraz2006*, we perturb the same number of 86 stimulation sites as used in the primate experiment. The site locations are randomly sampled from the unit locations. For *Afraz2015*, we run the same 40 optogenetic inactivation sessions as in the primate experiment with 17 of those sessions targeting face sites. Each suppression site is randomly sampled from the unit locations with a threshold of *d′ >* 1 for a site to count as “face selective.” For the muscimol benchmark, we follow the primate experiment and run 6 microinjections in face-selective sites and 6 in non-face-selective sites. For *Moeller2017*, a single primate experiment was run within and outside the face patch respectively. We deviate from the primate experimental paradigm in that we localize face patches with simulated fMRI recordings rather than biased sampling of locations. In the models, we can “record” every voxel (see *Model recordings*) and compute the per-voxel face-selectivity 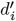 as the d-prime between face and non-face images:

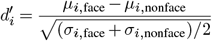

where *μ*_*i*,face_ and *μ*_*i*,nonface_ are neuron *i*’s mean response to a set of face and non-face images respectively, and *σ*_*i*,face_ and *σ*_*i*,nonface_ are neuron *i′s* variance values to face and non-face images respectively. For the stimulation inside the face patch we compute the purity center following Lee et al. (2020) with a radius of 1, i.e. the unit location where the sum of all voxels’ selectivities within 1mm is maximal. For the stimulation outside the face patch we randomly sample a unit location that is not part of a face patch with a threshold of *d′* = 0.85. For *Rajalingham2019*, we bootstrap the experiment 10 times with 10 sites each chosen uniformly from the 10 × 10mm artificial tissue sheet. Alignment scores to experimental data are computed for each bootstrap and then averaged.

#### Perturbation types

The different benchmarks use different types of perturbations with different parameters. *Afraz2006* micro-stimulate with a biphasic current pulse of 50 mA, a pulse rate of 200 Hz for 0.2 ms and a 0.1 ms interval. The stimulation onset was 100 ms after stimulus onset and lasted for 50 ms. For optogenetic suppression in *Afraz2015*, cortex is injected with a 6 *μL* solution at 0.1 *μL* per minute, containing AAV-8 carrying CAG-ARCHT. Per mL, the viral titter was 2 × 10^12^ infectious units. The laser pulse was then on for 200 ms with a total fiber output power of 12 mW. For muscimol suppression in *Afraz2015*, 1 *μL* of muscimol with 5 mg/*μL* was injected at a rate of 0.1 *μL* per minute. In *Moeller2017*, cortex was stimulated with a current pulse of 300 mA, a pulse rate of 150 Hz for 0.2 ms and a 0.1 ms interval. The stimulation lasted for 200 ms. For *Rajaling-ham2019*, 1 *μL* of muscimol with 5 mg/*μL* was injected at a rate of 0.1 *μL* per minute.

#### Behavioral measures

Most benchmarks use Δ% correct for measuring the effect of perturbations on behavior, except *Afraz2006* and *Rajalingham2019. Afraz2006* first fits the psychometric curve with a logistic function:

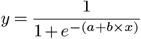

where *x* is a particular level of visual signal (Fig. 4), and *y* is the mean likelihood of choosing ‘face’ at that level. To determine the midpoint, we use the scipy ‘fsolve’ function with an initial guess of 0.5. The shift in behavior is then characterized as the shift in midpoints between non-stimulated and stimulated conditions. *Rajalingham2019* measures the behavioral effect in Δ*d′* on an object-level (‘O2’ Rajalingham et al., 2018). The *d′* value for objects *i* and *j* is computed as 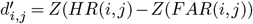 where the hit rate *HR* is the proportion of trials when subjects chose *i* when presented with images of object *i* against a distractor of object *j*, and the false alarms rate *FAR* is the proportion of trials when subject chose *i* when presented with images of object *j*.

### Base models

All models in this work are based on an AlexNet architecture with 224 × 224 image input trained on ImageNet 2012 for 100 epochs with stochastic gradient descent on a NVIDIA Titan X GPU, batch size 256, 0.01 initial learning rate decreased 10-fold every 40 epochs, 0.9 momentum, and 0.0001 weight decay. Model tissue is 10 × 10 mm. After training, behavioral choices are read out of fc6 with a logistic classifier, optimized with Newton’s method and *C* = 0.001.

#### TDANN via random arrangement

The model with random topography is trained without any additional parameters. Units’ x and y locations are assigned to tissue randomly by drawing from a uniform distribution. This model achieves a baseline accuracy of 57% top-1 performance on the ImageNet 2012 validation set.

#### TDANN via swapopt

The swapopt model starts from a fully trained model with random spatial layout, but post-hoc swaps unit locations to maximize spatial clustering with the same loss as for spatial co-training (Margalit et al., 2020). Specifically, we perform swapopt using ImageNet 2012 images in neighborhoods of size 6 mm, with 500 steps per neighbor-hood and 100,000,000 total steps.

#### TDANN via spatial co-training

To encourage spatial clustering, we co-train with a spatial loss in addition to the Ima-geNet categorization loss, following Lee et al. (2020). The model tissue spans 10 × 10 mm. The specific loss function is parameterized according to the response profile observed experimentally in primate IT (Lee et al., 2020):

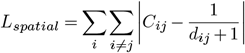

where *C*_*ij*_ is the correlation of response vectors between units *i* and *j* and *d*_*ij*_ is their Euclidean distance in millimeters in the tissue map. We apply this spatial loss to layers fc6 and fc7. The total loss is 0.1× categorization loss + 0.9 × the sum of spatial losses.

We test models trained from different seeds in this work. For visualizations, Fig. 4 is created from the seed 4 (because it looked the prettiest), and Fig. 5 and Table 1 from seed 0; both with a 44% top-1 accuracy. Note that while most seeds matched all qualitative results, not every single seed worked out. Anecdotally this seemed to be the result or a poor topography – studying such topographic model differences in more detail will be an interesting topic for future work. Differences between seeds are estimated in Fig. 6.

#### Alternatives

Alternative approaches to building a topographic model include a modification to the spatial loss used here (Margalit et al., 2023), the use of self-organizing feature maps (Kohonen, 1982), variational autoencoders (Keller and Welling, 2022), and a spatial connectivity cost on excitatory and inhibitory neural populations (Blauch et al., 2022), none of which were tested here.

### Ventral stream models

All models use *fc*6 as the ‘IT’ layer, and perform the behavioral readout from the same layer. All stimuli are assumed to be shown at the models’ full visual degrees of 8°, i.e. fill all pixels without padding.

#### Model recordings

To obtain fMRI recordings from the model, we smooth per-neuron spatial recordings with a Gaussian filter:

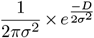

where 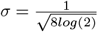 and *D* is used to convert from neuron locations to fMRI sampling locations with a resolution of 0.5mm.

To obtain Utah array recordings for 4 × 4mm electrode arrays with 10 × 10 electrodes and 0.4mm spacing, we use the same smoothing function with a resolution of 0.4mm.

#### Perturbation modules

Each perturbation module defines how the neural activity in a spatial profile changes as a result of a particular perturbation. The perturbed neural activity *N* ^*i*^ of a given layer is computed as the multiplicative delta with respect to the image-elicited activity *N* :

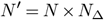

where *N*_Δ_ depends on the particular perturbation type.

### Muscimol

Suppressive muscimol injections at location *x*_*p*_, *y*_*p*_ are modeled with :

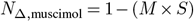

where the magnitude *M* is the injection amount in *μL* (here always *M* = 1 from 1*μL*) and the spatial spread *S* is the probability density function (pdf) of a multivariate normal distribution:

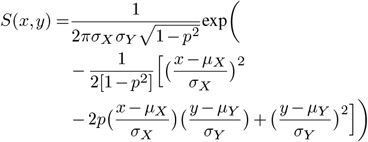

where the mean is the injection location, i.e. *μ*_*X*_ = *x*_*p*_ and *μ*_*Y*_ = *y*_*p*_, and *σ*_*X*_ = *σ*_*Y*_ = *p* = 1.17 modeled after experimental results in Arikan et al. (2002). *N*_Δ,muscimol_ is capped to not be smaller than 0.

### Optogenetics

Optogenetic inactivations are modeled the same way as muscimol suppression (above) with the pdf a multivariate normal distribution but the magnitude *M* is the fiber output power in *mW* and *σ* = 0.234 modeled after Chow et al. (2010) and Bernstein et al. (2008).

### Micro-stimulation

Activity increase following microstimulation is modeled after Kumaravelu et al. (2021):

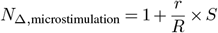

where *r* is the pulse rate in Hz, *R* = 30 is the assumed baseline firing rate following Ponce et al. (2019), and *S* is the spatial spread as an exponential fall off:

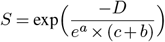

where *a* = 0.00505047368 and *b* = − 5.26062113 modeled after simulations in Kumaravelu et al. (2021), and *D* are the distances of unit locations *x, y* from the perturbation site *x*_*p*_, *y*_*p*_:

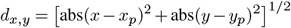

*N*_Δ,microstimulation_ is capped at a maximum of 2 which assumes a maximum firing rate of 2 × *R* = 60 Hz.

### Decoders

For all decoders that transform neural representations into behavioral task choices, we assume balanced training, i.e. we optimize decoders such that their performance on all classes should be identical. All categorization decoders use a linear (fully-connected) layer that transforms features into a softmax output corresponding to the number of classes. Features are first normalized to *μ* = 0 mean and *σ* = 1 standard deviation. The readout is then trained with cross-entropy and an Adam optimizer with a learning rate of 1*e* − 4 and 0.9 weight decay for 1, 000 epochs with a batch size of 100. For the *Moeller2017* same-different task, we implement a threshold decoder which computes the summed scalar distance between all features for the first and second stimulus. We then run binary search to determine the threshold that minimizes the difference between decoder performance on ‘same’ and ‘different’ trials. Note that this approach maximizes overall performance with near identical performances on each type of trial.

## References

Hamed Zivari Adab and Rufin Vogels. Perturbation of posterior inferior temporal cortical activity impairs coarse orientation discrimination. Cerebral Cortex, 26(9):3814–3827, 2016. doi: 10.1093/cercor/bhv178.

Arash Afraz, Edward S. Boyden, and James J. DiCarlo. Optogenetic and pharmacological suppression of spatial clusters of face neurons reveal their causal role in face gender discrimination. Proceedings of the National Academy of Sciences (PNAS), 112(21):6730–6735, 2015. doi: 10.1073/pnas.1423328112.

Seyed Reza Afraz, Roozbeh Kiani, and Hossein Esteky. Microstimulation of inferotemporal cortex influences face categorization. Nature, 442(7103):692–695, 2006. doi: 10.1038/nature04982.

Rasim Arikan, Nicquet M.J Blake, Joseph P Erinjeri, Thomas A Woolsey, Lisette Giraud, and Stephen M Highstein. A method to measure the effective spread of focally injected muscimol into the central nervous system with electrophysiology and light microscopy. Journal of Neuroscience Methods, 118:51–57, 2002. doi: 10.1016/S0165-0270(02)00143-7.

Mika Baba, Akiko Nishio, and Hidehiko Komatsu. Relationship Between the Activities of Gloss-Selective Neurons in the Macaque Inferior Temporal Cortex and the Gloss Discrimination Behavior of the Monkey. Cerebral Cortex Communications, 2(1):1–13, 2021. doi: 10.1093/TEXCOM/TGAB011.

Michael S. Beauchamp, Denise Oswalt, Ping Sun, Brett L. Foster, John F. Magnotti, Soroush Niketeghad, Nader Pouratian, William H. Bosking, and Daniel Yoshor. Dynamic Stimulation of Visual Cortex Produces Form Vision in Sighted and Blind Humans. Cell, 181(4):774–783, 2020. doi: 10.1016/j.cell.2020.04.033.

Jacob G Bernstein, Xue Han, Michael A Henninger, Emily Y Ko, Xiaofeng Qian, Giovanni Talei Franzesi, Jackie P. McConnell, Patrick Stern, Robert Desimone, and Edward S Boyden. Prosthetic systems for therapeutic optical activation and silencing of genetically targeted neurons. In Optical Interactions with Tissue and Cells XIX, volume 6854, page 68540H, 2008. doi: 10.1117/12.768798.

Nicholas M Blauch, Marlene Behrmann, and David C Plaut. A connectivity-constrained computational account of topographic organization in primate high-level visual cortex. Proceedings of the National Academy of Sciences (PNAS), 119(3), 2022. doi: 10.1073/pnas.2112566119.

Santiago A Cadena, George H Denfield, Edgar Y Walker, Leon A Gatys, Andreas S Tolias, Matthias Bethge, and Alexander S Ecker. Deep convolutional models improve predictions of macaque V1 responses to natural images. bioRxiv preprint, 2017. doi: 10.1101/201764.

Charles F. Cadieu, Ha Hong, Daniel L.K. Yamins, Nicolas Pinto, Diego Ardila, Ethan A. Solomon, Najib J. Majaj, and James J. DiCarlo. Deep Neural Networks Rival the Representation of Primate IT Cortex for Core Visual Object Recognition. PLoS Computational Biology, 10(12): 1–18, 2014. doi: 10.1371/journal.pcbi.1003963.

Charlotte Caucheteux and Jean-Rémi King. Brains and algorithms partially converge in natural language processing. Nature Communications Biology, 2022.

Ting Chen, Simon Kornblith, Mohammad Norouzi, and Geoffrey Hinton. A Simple Framework for Contrastive Learning of Visual Representations. arXiv preprint, 2020.

Brian Y. Chow, Xue Han, Allison S. Dobry, Xiaofeng Qian, Amy S. Chuong, Mingjie Li, Michael A. Henninger, Gabriel M. Belfort, Yingxi Lin, Patrick E. Monahan, and Edward S. Boyden. Highperformance genetically targetable optical neural silencing by light-driven proton pumps. Nature, 463(7277):98–102, 2010.

Joel Dapello, Tiago Marques, Martin Schrimpf, Franziska Geiger, David D. Cox, and James J. DiCarlo. Simulating a Primary Visual Cortex at the Front of CNNs Improves Robustness to Image Perturbations. In Neural Information Processing Systems (NeurIPS), 2020. doi: 10.1101/2020.06.16.154542.

Joel Dapello, Kohitij Kar, Martin Schrimpf, Robert Geary, Michael Ferguson, David D. Cox, and James J. DiCarlo. Aligning Model and Macaque Inferior Temporal Cortex Representations Improves Model-to-Human Behavioral Alignment and Adversarial Robustness. International Conference on Learning Representations (ICLR), 2023. doi: 10.1101/2022.07.01.498495.

Jia Deng, Wei Dong, Richard Socher, Li-Jia Li, Kai Li, and Li Fei-Fei. ImageNet: A large-scale hierarchical image database. In IEEE Conference on Computer Vision and Pattern Recognition, pages 248–255. IEEE, 2009. doi: 10.1109/CVPR.2009.5206848.

Adrien Doerig, Rowan P. Sommers, Katja Seeliger, Blake Richards, Jenann Ismael, Grace W. Lindsay, Konrad P. Kording, Talia Konkle, Marcel A. J. van Gerven, Nikolaus Kriegeskorte, and Tim C. Kietzmann. The neuroconnectionist research programme. Nature Reviews Neuroscience, 24(7):431–450, 2023. doi: 10.1038/s41583-023-00705-w.

Eduardo Fernández, Arantxa Alfaro, Cristina Soto-Sánchez, Pablo Gonzalez-Lopez, Antonio M. Lozano, Sebastian Peña, Maria Dolores Grima, Alfonso Rodil, Bernardeta Gómez, Xing Chen, Pieter R. Roelfsema, John D. Rolston, Tyler S. Davis, and Richard A. Normann. Visual percepts evoked with an intracortical 96-channel microelectrode array inserted in human occipital cortex. Journal of Clinical Investigation, 131(23), 2021. doi: 10.1172/jci151331.

Franziska Geiger, Martin Schrimpf, Tiago Marques, and James J Dicarlo. Wiring Up Vision : Minimizing Supervised Synaptic Updates Needed to Produce a Primate Ventral Stream. International Conference on Learning Representations (ICLR), 2022. doi: 10.1101/2020.06.08.140111.

Mark H. Histed, Vincent Bonin, and R. Clay Reid. Direct Activation of Sparse, Distributed Populations of Cortical Neurons by Electrical Microstimulation. Neuron, 63(4):508–522, 2009. doi: 10.1016/J.NEURON.2009.07.016.

Gary B Huang, Manu Ramesh, Tamara Berg, and Erik Learned-Miller. Labeled Faces in the Wild: A Database for Studying Face Recognition in Unconstrained Environments. Technical report, University of Massachusetts, Amherst, 2007.

Mehrdad Jazayeri, Zachary Lindbloom-Brown, and Gregory D. Horwitz. Saccadic eye movements evoked by optogenetic activation of primate V1. Nature Neuroscience, 15(10):1368–1370, 2012. doi: 10.1038/nn.3210.

Keisuke Kawasaki and David L. Sheinberg. Learning to recognize visual objects with microstimulation in inferior temporal cortex. Journal of Neurophysiology, 100(1):197–211, 2008. doi: 10.1152/jn.90247.2008.

Alexander J.E. Kell, Daniel L.K. Yamins, Erica N. Shook, Sam V. Norman-Haignere, and Josh H. McDermott. A Task-Optimized Neural Network Replicates Human Auditory Behavior, Predicts Brain Responses, and Reveals a Cortical Processing Hierarchy. Neuron, 98(3):630–644, 2018. doi: 10.1016/j.neuron.2018.03.044.

T. Anderson Keller and Max Welling. Topographic VAEs learn Equivariant Capsules. arXiv, 2022. doi: 10.48550/arXiv.2109.01394.

Seyed Mahdi Khaligh-Razavi and Nikolaus Kriegeskorte. Deep Supervised, but Not Unsupervised, Models May Explain IT Cortical Representation. PLoS Computational Biology, 10(11), 2014. doi: 10.1371/journal.pcbi.1003915.

Teuvo Kohonen. Self-organized formation of topologically correct feature maps. Biological Cybernetics, 43(1):59–69, 1982. doi: 10.1007/BF00337288.

Jonas Kubilius, Martin Schrimpf, Ha Hong, Najib J. Majaj, Rishi Rajalingham, Elias B. Issa, Kohitij Kar, Pouya Bashivan, Jonathan Prescott-Roy, Kailyn Schmidt, Aran Nayebi, Daniel Bear, Daniel L. K. Yamins, and James J. DiCarlo. Brain-Like Object Recognition with High-Performing Shallow Recurrent ANNs. In Neural Information Processing Systems (NeurIPS), pages 12785—-12796, 2019.

Karthik Kumaravelu, Joseph Sombeck, Lee E Miller, Sliman J Bensmaia, and Warren M Grill. Stoney vs. Histed: Quantifying the Spatial Effects of Intracortical Microstimulation. bioRxiv preprint, 2021. doi: 10.1101/2021.08.12.456091.

Hyodong Lee, Eshed Margalit, Kamila M Jozwik, Michael A Cohen, Nancy Kanwisher, Daniel L K Yamins, and James J Dicarlo. Topographic deep artificial neural networks reproduce the hallmarks of the primate inferior temporal cortex face processing network. bioRxiv preprint, page 2020.07.09.185116, 2020. doi: 10.1101/2020.07.09.185116.

Najib J Majaj, Ha Hong, Ethan A Solomon, and James J DiCarlo. Simple Learned Weighted Sums of Inferior Temporal Neuronal Firing Rates Accurately Predict Human Core Object Recognition Performance. Journal of Neuroscience, 35(39):13402–13418, 2015. doi: 10.1523/JNEUROSCI.5181-14.2015.

Eshed Margalit, Hyodong Lee, Tiago Marques, James J. DiCarlo, and Daniel Yamins. Correlation-based spatial layout of deep neural network features gen-erates ventral stream topography. In Computational and Systems Neuroscience (CoSyNe), pages 242—-243, 2020.

Eshed Margalit, Hyodong Lee, Dawn Finzi, James J. DiCarlo, Kalanit Grill-Spector, and Daniel L. K. Yamins. A Unifying Principle for the Functional Organization of Visual Cortex. arXiv, 2023. doi: 10.1101/2023.05.18.541361.

Sebastian Moeller, Trinity Crapse, L. Chang, and Doris Y. Tsao. The effect of face patch microstimulation on perception of faces and objects. Nature Neuroscience, 20(5):743–752, 2017. doi: 10.1038/nn.4527.

Aran Nayebi, Alexander Attinger, Malcolm G Campbell, Kiah Hardcastle, Isabel I.C. Low, Caitlin S Mallory, Gabriel C Mel, Ben Sorscher, Alex H Williams, Surya Ganguli, Lisa M Giocomo, and Daniel L.K. Yamins. Explaining heterogeneity in medial entorhinal cortex with task-driven neural networks. In Neural Information Processing Systems (NeurIPS), volume 34, 2021.

Carlos R. Ponce, Will Xiao, Peter F. Schade, Till S. Hartmann, Gabriel Kreiman, and Margaret S. Livingstone. Evolving Images for Visual Neurons Using a Deep Generative Network Reveals Coding Principles and Neuronal Preferences. Cell, 177(4):999–1009, 2019. doi: 10.1016/j.cell.2019.04.005.

Rishi Rajalingham and James J. DiCarlo. Reversible Inactivation of Different Millimeter-Scale Regions of Primate IT Results in Different Patterns of Core Object Recognition Deficits. Neuron, 102(2):493–505, 2019. doi: 10.1016/j.neuron.2019.02.001.

Rishi Rajalingham, Elias B. Issa, Pouya Bashivan, Kohitij Kar, Kailyn Schmidt, and James J DiCarlo. Large-Scale, High-Resolution Comparison of the Core Visual Object Recognition Behavior of Humans, Monkeys, and State-of-the-Art Deep Artificial Neural Networks. The Journal of Neuroscience, 38(33):7255–7269, 2018. doi: 10.1523/JNEUROSCI.0388-18.2018.

Srivatsun Sadagopan, Wilbert Zarco, and Winrich A. Freiwald. A causal relationship between face-patch activity and face-detection behavior. eLife, 6, 2017. doi: 10.7554/eLife.18558.

Mark R. Saddler, Ray Gonzalez, and Josh H. McDermott. Deep neural network models reveal interplay of peripheral coding and stimulus statistics in pitch perception. Nature Communications, 12(1):7278, 2021. doi: 10.1038/s41467-021-27366-6.

Martin Schrimpf, Jonas Kubilius, Ha Hong, Najib J. Majaj, Rishi Rajalingham, Elias B. Issa, Kohitij Kar, Pouya Bashivan, Jonathan Prescott-Roy, Kailyn Schmidt, Daniel L. K. Yamins, and James J. DiCarlo. Brain-Score: Which Artificial Neural Network for Object Recognition is most Brain-Like? bioRxiv, 2018. doi: 10.1101/407007.

Martin Schrimpf, Jonas Kubilius, Michael J Lee, N. Apurva Ratan Murty, Robert Ajemian, and James J. DiCarlo. Integrative Benchmarking to Advance Neurally Mechanistic Models of Human Intelligence. Neuron, 2020. doi: 10.1016/j.neuron.2020.07.040.

Martin Schrimpf, Idan Asher Blank, Greta Tuckute, Carina Kauf, Eghbal A. Hosseini, Nancy Kanwisher, Joshua B. Tenenbaum, and Evelina Fedorenko. The neural architecture of language: Integrative modeling converges on predictive processing. Proceedings of the National Academy of Sciences (PNAS), 118(45):e2105646118, 2021. doi: 10.1073/pnas.2105646118.

S D Stoney, W D Thompson, and H Asanuma. Excitation of Pyramidal Tract Cells by Intracortical Microstimulation: Effective Extent of Stimulating Current. Journal of Neurophysiology, 31(5): 659–69, 1968. doi: 10.1152/jn.1968.31.5.659.

Mariya Toneva and Leila Wehbe. Interpreting and improving natural-language processing (in machines) with natural language-processing (in the brain). In Advances in Neural Information Processing Systems, volume 32, pages 14954–14964, 2019.

Daniel L K Yamins and James J DiCarlo. Using goal-driven deep learning models to understand sensory cortex. Nature Neuroscience, 19(3):356–365, 2016. doi: 10.1038/nn.4244.

Daniel L. K. Yamins, Ha Hong, Charles F. Cadieu, Ethan A. Solomon, Darren Seibert, and James J. DiCarlo. Performance-optimized hierarchical models predict neural responses in higher visual cortex. Proceedings of the National Academy of Sciences (PNAS), 111(23): 8619–8624, 2014. doi: 10.1073/pnas.1403112111.

Chengxu Zhuang, Jonas Kubilius, Mitra JZ Hartmann, and Daniel L. Yamins. Toward Goal-Driven Neural Network Models for the Rodent Whisker-Trigeminal System. In Neural Information Processing Systems (NIPS), pages 2555–2565, 2017.

Chengxu Zhuang, Siming Yan, Aran Nayebi, Martin Schrimpf, Michael C. Frank, James J. DiCarlo, and Daniel L.K. Yamins. Unsupervised neural network models of the ventral visual stream. Proceedings of the National Academy of Sciences (PNAS), 118(3), 2021. doi: 10.1073/pnas.2014196118.

